# SCOPE: Revealing Hidden Mechanisms in Phenotypic Screens Through Target and Pathway Enrichment

**DOI:** 10.1101/2025.07.11.664427

**Authors:** Abhijeet Kapoor, Keith Kelleher, Suzanne Underhill, Sankalp Jain, Brandon K Harvey, Mark J. Henderson

## Abstract

Phenotypic screening enables discovery of small molecules without requiring predefined targets, but mechanistic interpretation remains challenging due to polypharmacology and pathway complexity. We developed SCOPE (Screening Compound Ontology for Pathway Enrichment), a KNIME-based computational framework that resolves the molecular drivers of phenotypic activity by linking compound-level screening data to annotated targets and pathways. SCOPE integrates multi-source target annotations and performs statistical enrichment to identify shared mechanisms of action. Applied to a high-throughput screen for modulators of ER-stress induced secretion of endoplasmic reticulum (ER) resident proteins, a process known as exodosis, SCOPE identified calcium signaling as the most enriched KEGG pathway without prior biological context. Target enrichment revealed G protein-coupled receptors (GPCRs) involved in inositol 1,4,5-trisphosphate receptors (IP3Rs)-mediated signaling, with widespread antagonism among hit compounds implicating this pathway in the regulation of exodosis. Notably, SCOPE uncovered a novel role for the histamine receptor HRH1, which was validated by RNAi knockdown and pharmacological inhibition, implicating HRH1 as a potential therapeutic target in ER stress-related disorders. These results highlight SCOPE’s potential to deconvolute phenotypic screens and uncover actionable mechanisms in complex cellular systems.

---

In recent years, phenotypic screening has gained momentum as a practical and fruitful strategy in early drug discovery. While target-based approaches have contributed to many therapeutic successes, they often require detailed molecular understanding of disease mechanisms and may fail to capture the biological complexity underlying multifactorial disorders^1^. In contrast, phenotypic assays evaluate the functional effects of small molecules in intact cellular systems, enabling the discovery of active compounds without prior assumptions about their molecular targets^2^. By capturing biologically meaningful, system-level responses, phenotypic screening is particularly well suited to interrogate complex processes involving polypharmacology, pathway crosstalk, and poorly characterized molecular drivers. Phenotypic screeing has been especially valuable in areas such as cancer, neurodegeneration, and metabolic disorders, where traditional reductionist methods may not sufficiently capture therapeutically relevant biology^3^. Advances in high-throughput screening formats, assay multiplexing, automated microscopy, and integrative data analysis have further enhanced the scalability and precision of phenotypic approaches. These developments have enabled researchers to systematically interrogate thousands of compounds across disease-relevant cellular models, accelerating both mechanism-agnostic drug discovery and the repurposing of approved compounds^4^.

One such biological process that has been used for phenotypic interrogation is the regulation of the endoplasmic reticulum (ER) proteome, which governs essential functions such as protein folding, post-translational modification, calcium signaling, and stress response activation^5–9^. The ER plays a critical role in maintaining protein homeostasis (proteostasis), and disturbances to its environment, particularly disruptions in calcium balance, can trigger the accumulation of misfolded proteins and initiate the unfolded protein response (UPR). In extreme cases, these disruptions lead to a recently described phenomenon known as exodosis: the stress-induced secretion of ER-resident proteins into the extracellular space^6,10^. This non-canonical release bypasses KDEL-dependent retention mechanisms and includes the export of molecular chaperones and folding enzymes, which are typically confined to the ER lumen. Exodosis has been increasingly recognized as a signature of ER dysfunction and has been observed in models of neurodegeneration, hyperthermia, ischemia-reperfusion injury, diabetes, and certain cancers, conditions often characterized by persistent ER stress and calcium dysregulation^6,11–13^. Despite its growing clinical relevance, the mechanisms driving exodosis remain poorly defined, and no therapies currently exist that directly target or modulate this secretory response. This presents a critical opportunity for phenotype-driven approaches to identify novel small-molecule modulators and uncover the biological pathways involved in maintaining ER proteome integrity under stress.

In a prior study, we created a *Gaussia* luciferase-based secreted ER calcium-modulated protein (GLuc SERCaMP) reporter, which enabled a high-throughput phenotypic screen to identify small molecules that inhibit calcium depletion-induced exodosis^6^. Among the top characterized hits was bromocriptine, a dopamine receptor agonist with established clinical use but a poorly understood mechanism in the context of ER stress. While this work confirmed the feasibility of phenotypic screening to uncover modulators of ER proteostasis, mechanistic interpretation of the full set of hit compounds was obscured by their chemical diversity and potential for polypharmacology. This challenge remains common in phenotypic screens, where chemically diverse and often polypharmacological hit profiles complicate efforts to map biological activity to specific molecular targets or signaling pathways. Consequently, there is a critical need for computational methods that can translate a list of small molecules identified in a phenotypic assay into biologically interpretable insights by systematically identifying shared molecular features and pathway-level convergence.

To address the challenge of mechanistic interpretation following phenotypic screening, a number of computational tools have been developed to support enrichment analysis and post hoc annotation of compound sets. Drug Set Enrichment Analysis (DSEA)^14^ leverages transcriptional perturbation signatures from the Connectivity Map (CMap) to detect shared pathways modulated by groups of phenotypically similar compounds. While informative, DSEA relies primarily on gene expression-based associations and is constrained by the limited chemical and pathway coverage of CMap data. Other tools such as DrugPattern^15^, Drugmonizome^16^, and CSgator^17^ perform drug set enrichment analysis using curated annotations for drug-target interactions, pathways, diseases, and side effects. Path4Drug^18^ integrates compound-target mappings with Reactome^19^ pathway annotations in a tissue-specific context, with a primary focus on drug safety and toxicity assessment. While these tools offer a relatively broader coverage of chemical space, they typically perform enrichment against a fixed universe of reference compounds, which may not reflect the specific chemical space represented in a given phenotypic screen.

To overcome these limitations, we developed SCOPE (Screening Compound Ontology for Pathway Enrichment), a computational enrichment pipeline that enables pathway deconvolution tailored to the phenotypic screening dataset. SCOPE integrates curated compound-target interaction data from publicly available sources including ChEMBL^20^, IUPHAR^21^, PubChem^22^, PharmGKB^23^, and DrugBank^24^ as well as any user supplied annotations. The compound-target mapping is performed for both the hit compounds and for the entire screening library, enabling a compound-target association matrix that reflects the experimental context. Enrichment analysis is then performed to identify targets overrepresented among hit compounds compared to this empirically defined background. Subsequently, the enriched targets are subjected to functional enrichment using STRING^25^ pathway annotations to identify associated biological processes and signaling cascades.

We applied SCOPE to a previously published phenotypic screen of 10,124 small molecules, from the NCATS bioactive collections, which identified compounds that inhibited ER calcium depletion-induced exodosis^6^. Our prior analyses positioned bromocriptine as a polypharmacological agent acting through non-canonical signaling pathways. SCOPE identified enrichment of GPCR targets linked to IP3R-mediated calcium signaling, implicating this axis in the exodosis response. It also uncovered a previously unrecognized role for the histamine receptor HRH1 in exodosis, which we validated experimentally through RNAi knockdown and chemical antagonism with selective small molecule inhibitors. These findings illustrate how pathway-level deconvolution using SCOPE can provide mechanistic insights into phenotypic hits, reveal novel therapeutic targets, and support rational drug development.

## Results

### Target enrichment of screening hits identified as modulators of exodosis

To identify small-molecule modulators of ER exodosis, we previously caried out quantitative HTS of 10,124 compounds which led to the identification of 185 compounds that reduced exodosis, which were interpreted to function through diverse mechanisms of action^6^. However, the pharmacologically diverse array of hits alone did not provide a clear picture of the underlying targets or mechanism of actions. Here, we took a data-centric approach, using the 185 compounds hit list together with the entire screening collections as inputs to our SCOPE workflow (Fig. 1; see methods for details) to systematically connect hits from the phenotypic screen to an intricate web of biological pathways.

**Figure 1.**
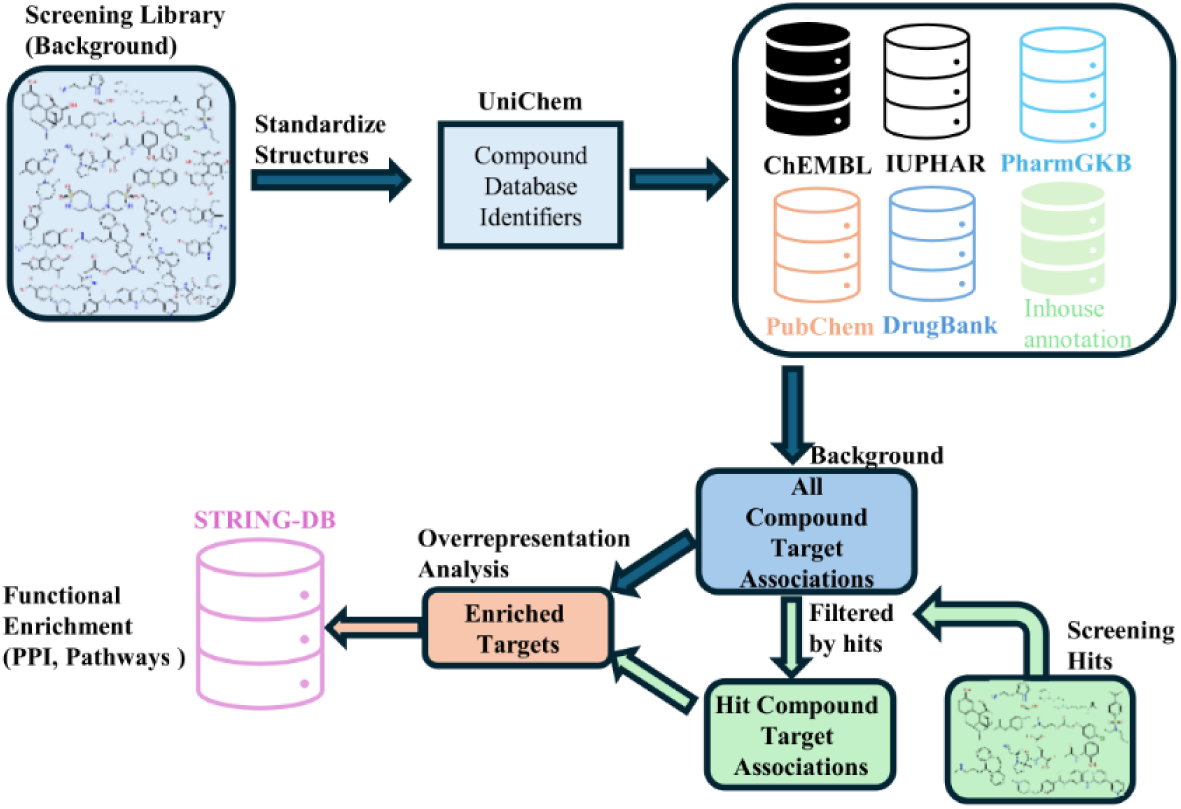
Schematic representation of the SCOPE workflow.

Running the SCOPE workflow on the entire screening collection (10,124 compounds) resulted in compound-target profiles for 6,829 compounds while no target association data was retrieved for remaining 3,295 compounds. A total of 44,771 compound-target pairs were obtained for the 6,829 compounds resulting in association with 2,702 unique targets. For the 185 compounds in the hit list, compound-target profiles were obtained for 143 hits, with 1331 compound-target pairs resulting in 392 unique targets. A target enrichment analysis comparing the compound-target profiles of the hit list with those of the background identified 26 targets enriched at 5% FDR and 70 targets enriched at 5% p-value (Supplementary Table 1). A functional enrichment analysis of the entire set of 392 mapped targets, conducted using the STRING database, identified 170 enriched KEGG pathways, with various metabolism- and addiction-related pathways among the top ten, while the calcium signaling pathway ranked much lower (Fig. 2A). In contrast, enrichment analysis using the 26 enriched targets identified by SCOPE (FDR < 5%) revealed only 11 enriched KEGG pathways, with calcium signaling emerging as the top hit (Fig. 2B). Notably, an analysis of the 70 enriched targets, identified by a 5% p-value cutoff, still highlighted the calcium signaling pathway as the topmost among 17 enriched KEGG pathways (Fig. 2C), further validating that SCOPE effectively filters background noise and isolates biologically relevant signals. These results underscore the ability of SCOPE to highlight specific pathway, i.e. calcium signaling a key biological pathway of interest without prior knowledge of the screening hit’s mode of action.

**Figure 2.**
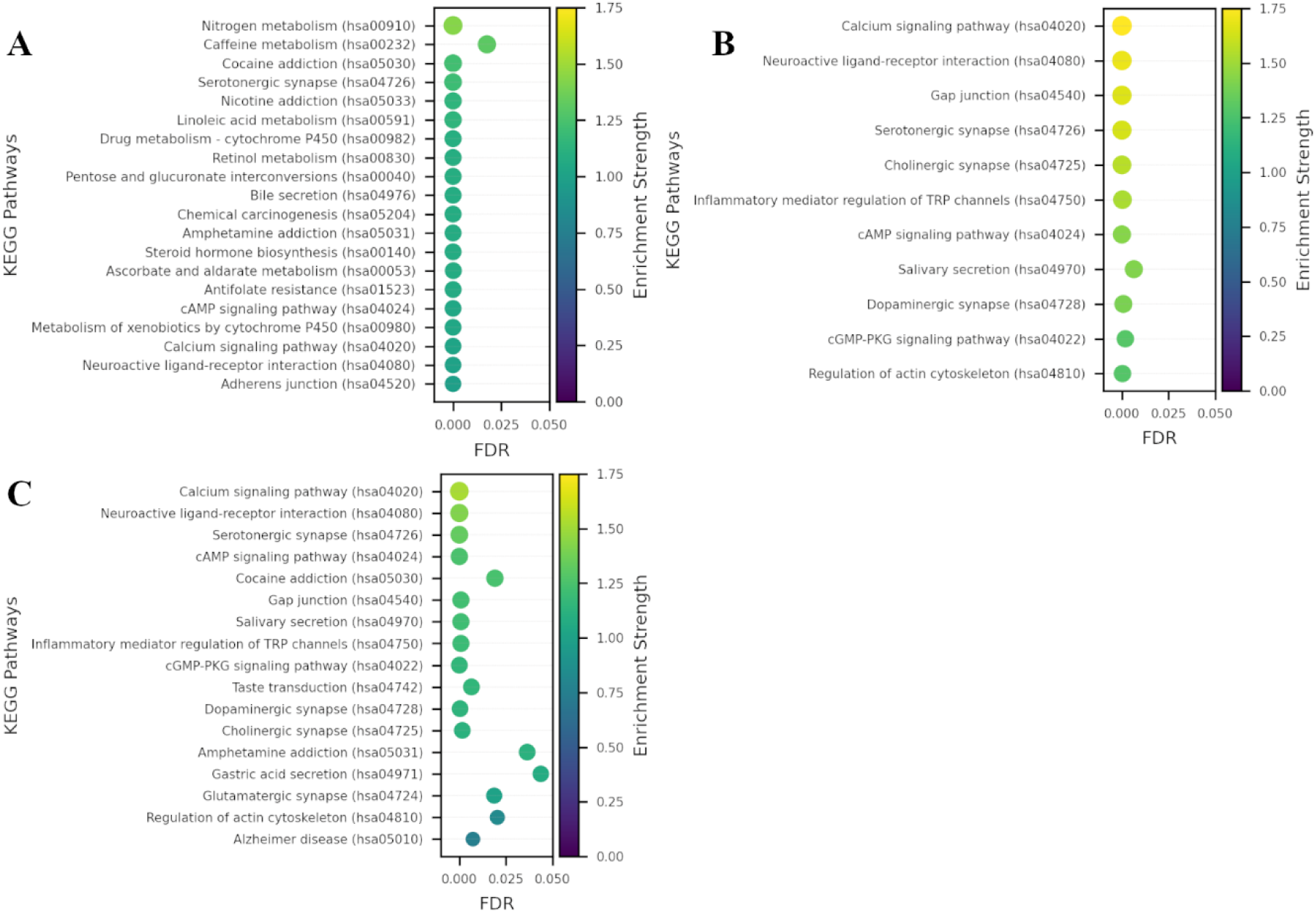
Enriched KEGG pathways identified by STRING functional enrichment analysis using **A)** all 392 mapped targets for 185 screening hits, **B)** 26 enriched targets identified by 5% FDR cutoff after target enrichment analysis, and **C)** 70 enriched targets identified by 5% p-value cutoff after target enrichment analysis. The pathways are ranked by the enrichment strength metric from STRING, which quantifies the magnitude of enrichment as the base 10 logarithm of the ratio between the observed and expected number of proteins annotated with a given term.

### Polypharmacology and GPCR-mediated regulation of ER exodosis

In our previous study, we prioritized five FDA-approved hits from the screen for further investigation. Dextromethorphan, bromocriptine, dantrolene, verapamil, and diltiazem were selected, due to their known roles in calcium modulation, with particular focus on characterizing bromocriptine^6^. Although bromocriptine is a dopamine D2 receptor (DRD2) agonist, our data suggested that bromocriptine’s anti-exodosis activity was unrelated to its dopaminergic activity leading us to suggest a non-dopamine receptor type GPCR, or more likely a distinct yet unknown target, as mediating bromocriptine’s ER proteome-stabilizing effect^6^. SCOPE analysis revealed that bromocriptine exhibits a more nuanced polypharmacology wherein it acts as agonists for a subset of dopamine and serotonin receptors while antagonizing the activity of adrenergic and certain other dopamine and serotonin receptors (Supplementary Table 2). The analysis brings forward another possible explanation for bromocriptine’s effect on exodosis, in which a complex interplay of GPCR actions are driving the effect. This possibility was overlooked in our earlier interpretations of the data, as our efforts primarily focused on evaluating dopamine receptor modulation.

Among the 26 enriched targets, we observed prominent overrepresentation of GPCRs—particularly muscarinic (CHRM1-5), serotonin (HTR1A, HTR2A, HTR2B, HTR2C, HTR3A, HTR6), adrenergic (ADRA1A, ADRA1D, ADRA2A, ADRA2B), histamine (HRH1, HRH2), and dopamine (DRD1-3, DRD5) receptors (Fig. 3). These GPCRs couple to G_q_ proteins and activate the phospholipase C (PLC) and the IP3R signaling pathway resulting in the IP3 receptor mediated release of Ca^2+^ from the ER^26–28^. Based on this shared signaling axis, we hypothesized that hit compounds may attenuate exodosis by antagonizing the IP3R pathway. To further explore this hypothesis, we first analyzed the compound-target activity data for the hits that associated with 26 enriched targets to classify the action of compound on targets (see methods for details). As shown in Fig. 4, 55 out of 185 hits associated with 26 enriched targets with several of the hits exerting broad inhibitory effects, potentially functioning as pan-antagonists across multiple GPCRs (compound action ‘inhibitor’ highlighted in red circles in Fig. 4) while for some other broadly acting compounds the mode of action could not be automatically assigned based on activity data alone (those labelled ‘inconclusive’). Notably, a subset of hits displayed preferential inhibition of one GPCR among the 26 enriched targets, providing an opportunity to dissect more discrete regulatory roles for these targets. These included compounds such as NCGC00016366 (azacyclonol) which targets histamine receptor (HRH1), NCGC00024378 (lobeline) which targets HTR2A, NCGC00024788 (2-CMDO) which targets DRD2, NCGC00024880 (AR-C239) that inhibits ADRA1A, NCGC00025328 (SB-258585) that inhibits HTR6, and NCGC00025335 (esreboxetine) which targets SLC6A2. Notably, in addition to azacyclonol, several FDA-approved selective antihistamines also appeared in this list despite showing some off-target activity (Fig. 4). These included ketotifen (NCGC00015580), cyclizine (NCGC00016421), clemastine (NCGC00016710), desloratadine (NCGC00159325), and chlorcyclizine (NCGC00179384). These findings prompted us to further investigate the role of histamine receptor signaling in modulating the exodosis response.

**Figure 3.**
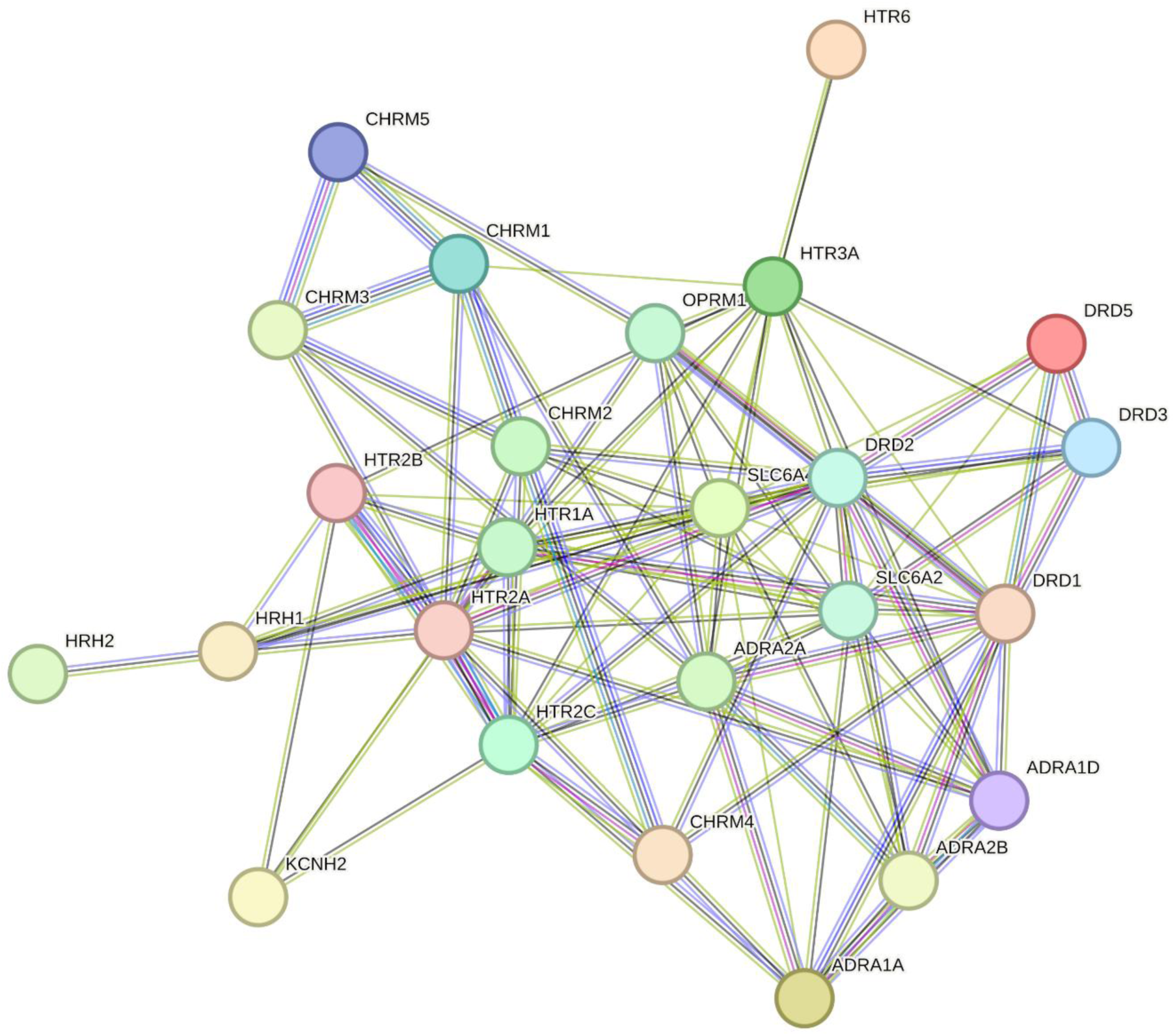
Protein-protein interaction network from STRING for the 26 enriched targets identified by 5% FDR cutoff after target enrichment analysis.

**Figure 4.**
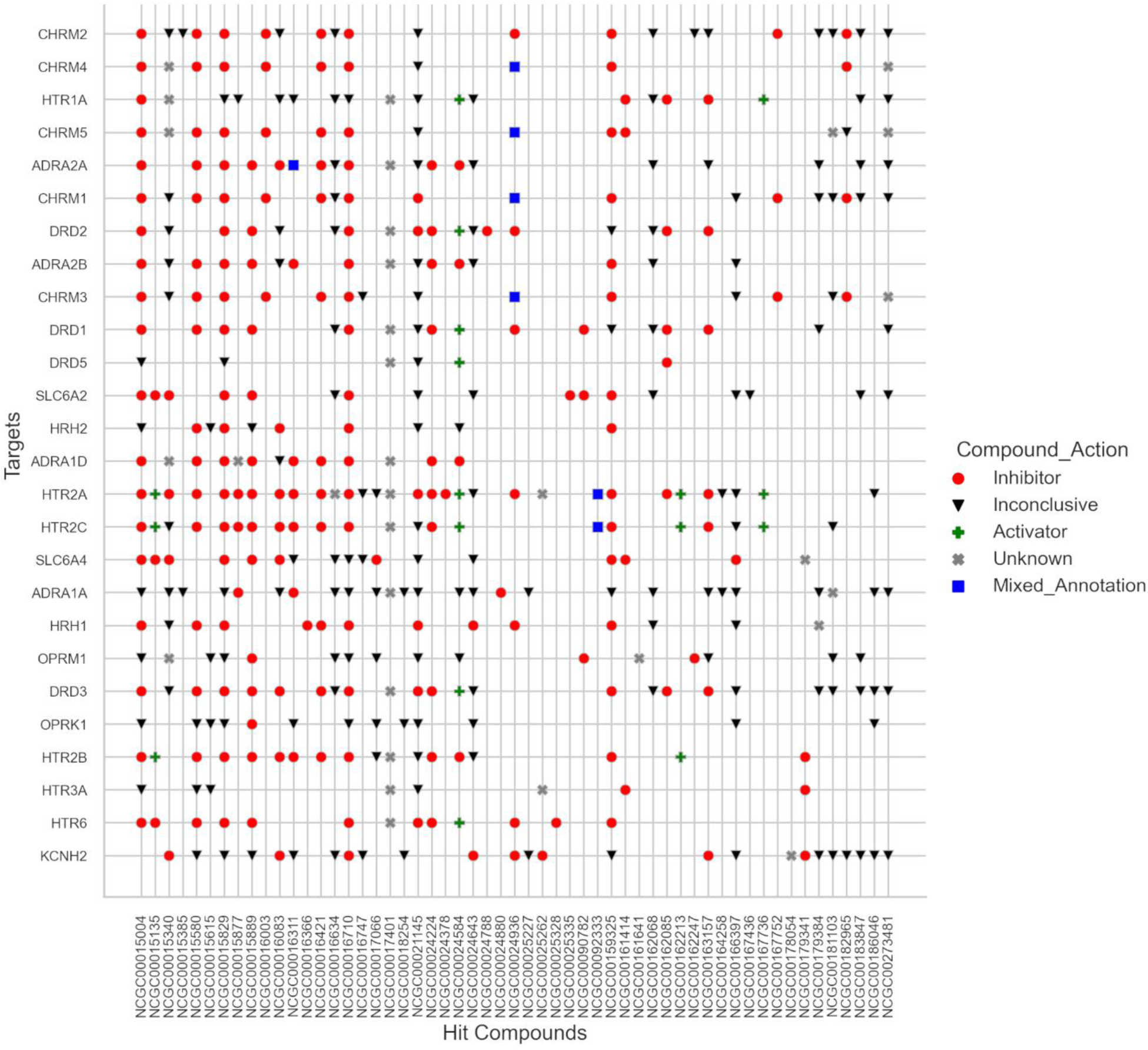
Classification of 55 hit compounds (NCGC IDs) action associated with the 26 enriched targets based on reported activity annotations. Compounds were classified as inhibitors if annotated with K_i_, pK_i_, INH, or inhibitory concentrations (e.g., IC_50_, IC_20_), and as activators if associated with EC values (e.g., EC_50_, EC_20_). Metrics unrelated to functional modulation (e.g., K_d_, K_b_, K_m_, K_a_) were labeled inconclusive. Compounds annotated as both activators and inhibitors for the same target were marked as mixed. Associations from curated sources (e.g., DrugBank, PharmGKB) without specific activity values were labeled unknown.

### Histamine receptor silencing attenuates exodosis

To investigate the role of histamine signaling in exodosis, we utilized the previously described GLuc SERCaMP assay^6^ to monitor ER calcium release following thapsigargin-induced ER stress in SH-SY5Y human neuroblastoma cells. We tested three selective histamine receptor antagonists: rupatadine^29^ (pIC_50_ 8.4), azelastine^30^ (pK_i_ 8.9; FDA approved), and clemastine^31^ (pK_i_ 10.3; FDA approved), and observed that all three compounds significantly reduced GLuc SERCaMP signal relative to control suggesting anti-exodosis activity (Fig. 5A). In contrast, selective activation of HRH1 using two selective agonists: NCGC00186016-01 (*N-*methylhistaprodifen)^32^ and NCGC00159581-01 (2-pyridylethylamine)^33^, resulted in an increase in GLuc SERCaMP signal suggesting an increase in exodosis response (Fig. 5B). To further confirm the role of histamine receptor in regulating the exodosis response, we examined expression of the histamine receptor paralogs in SH-SY5Y cells, which revealed highest expression of HRH1, with near undetectable levels of HRH2-4 (Fig 5C). Next, we performed an RNAi-mediated knockdown of HRH1 in SH-SY5Y GLuc SERCaMP reporter cells. Following HRH1 knockdown, we observed a significant reduction in SERCaMP secretion without significant loss in cell viability (Fig. 5D), further validating that HRH1 contributes to the regulation of ER calcium dynamics and may play a role in mediating the exodosis phenotype.

**Figure 5.**
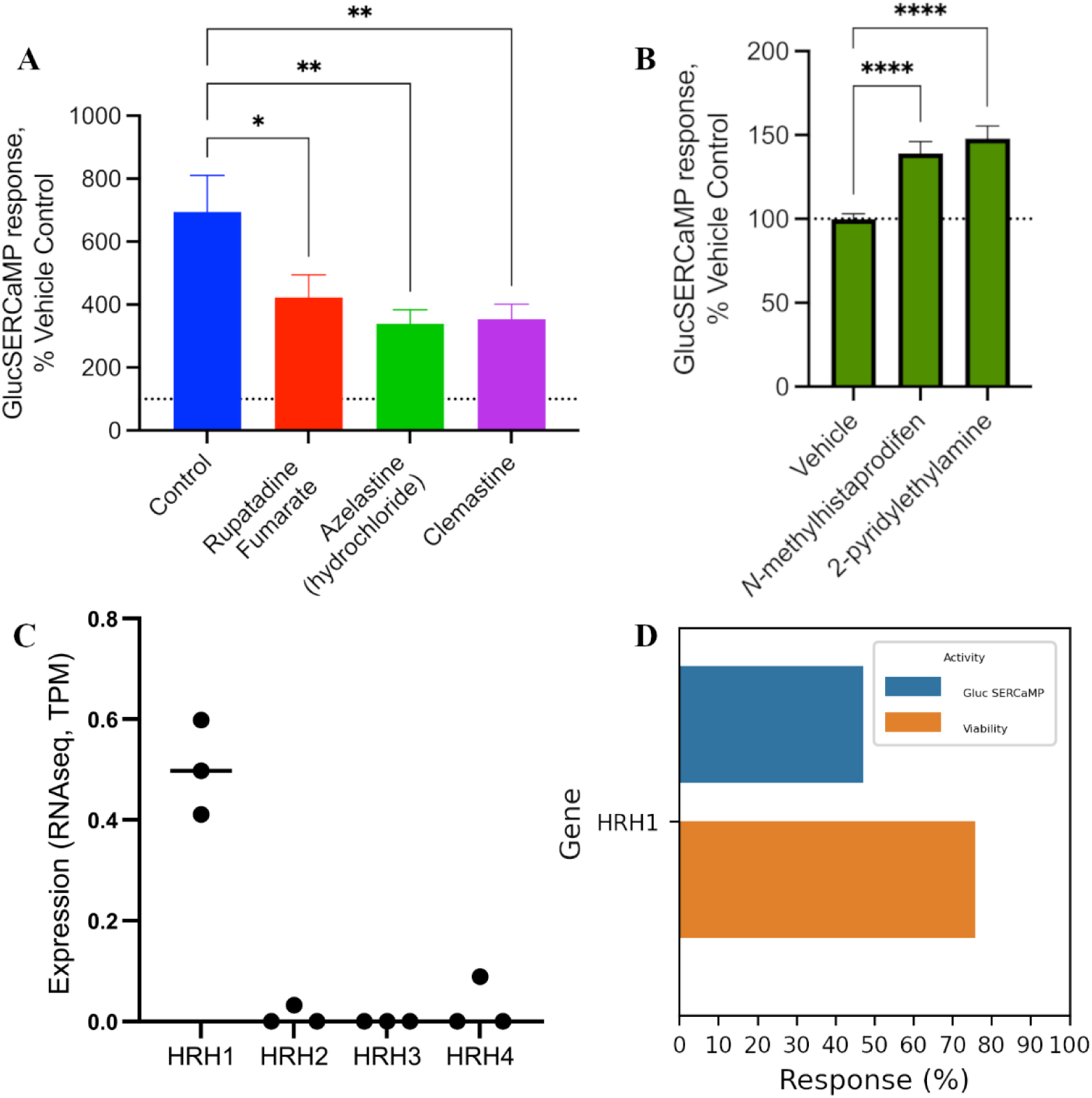
Effect of histamine receptor 1 (HRH1) on exodosis response. **A)** Stable GLuc SERCaMP SH-SY5Y cells were pre-treated with 10 uM of one of three selective H1 antagonists (rupatadine, azelastine, and clemastine) for 16 hours. Subsequently, they were treated with 100 nM thapsigargin for four hours. Data here represents the results of three different experiments all normalized to their own Vehicle control condition. (*<0.05, **<0.01, ****<0.0001). **B)** Two different selective H1 receptor agonists (*N-*methylhistaprodifen and 2-pyridylethylamine) were applied to stable GLuc SERCaMP SH-SY5Y cells for 16 hours. (10 μM, *<0.05, **<0.01, ****<0.0001). **C)** RNAseq data examining the expression of the histamine receptor paralogs in SH-SY5Y cells. **D)** GLuc SERCaMP and viability for cells with RNAi-mediated knockdown of HRH1. Values reported correspond to the median of three independent SERCaMP and viability runs.

## Discussion

The re-emergence of phenotypic screening in early drug discovery has highlighted its value in probing complex biological processes without prior assumptions about molecular targets, particularly in multifactorial disorders where traditional target-based approaches may fall short. However, interpreting the results of such screens remains challenging due to the pharmacological and structural diversity of hits, the polypharmacology of small molecules, and the lack of mechanistic context linking compounds to pathways. Manual analysis of large (or even moderately sized) hit lists to gain biological insights can be a cumbersome task, often leading investigators to focus on a much smaller subset of hits and prioritize results that align with prior hypotheses, thereby missing unexpected or novel relationships embedded in the dataset. To address these challenges and uncover relationships with intricate biological networks, we developed the Screening Compound Ontology for Pathway Enrichment (SCOPE) platform, an extensible KNIME-based pipeline that integrates compound-target annotation with pathway-level analysis to systematically infer mechanisms of action underlying phenotypic screening hits. SCOPE leverages the specific chemical and target space represented in a given screening library to identify protein targets statistically overrepresented among hit compounds from a phenotypic screen, followed by functional enrichment analysis to identify associated biological processes. This two-step approach, looking at target-level enrichment followed by pathway-level inference, enables a more accurate and biologically relevant interpretation of phenotypic screening results, improving upon existing tools and provides deeper mechanistic insight into the molecular drivers of phenotypic activity.

In this study, we applied SCOPE to a target-agnostic high-throughput screen for stabilizers of the ER resident proteome^6^, yielding significant insights into the mechanisms underlying compound activity. The hit compounds from this screen reduced secretion of GLuc SERCaMP, a reporter of exodosis signifying calcium depletion-induced ER stress response. While a broad functional enrichment analysis of all 392 mapped targets from the hit list identified numerous pathways, including metabolism- and addiction-related ones, SCOPE’s target enrichment analysis identified 26 enriched targets, which sharply focused the biological signal, revealing calcium signaling as the top enriched KEGG pathway. This underscores SCOPE’s effectiveness in filtering background noise and isolating biologically relevant signals without prior knowledge of the compound’s mode of action, a key advantage over simpler target mapping approaches. By applying a focused enrichment strategy, SCOPE highlights specific molecular regulators likely to mediate ER exodosis, revealing putative mechanisms that would be difficult to discern using all mapped targets alone.

The prominent representation of GPCRs among the 26 enriched targets, including muscarinic, serotonin, adrenergic, histamine, and dopamine receptors, is particularly noteworthy and suggests that these receptors may serve as signaling hubs whose pharmacological modulation alters the cellular response to ER calcium depletion. These GPCRs are known to couple to G_q_ proteins, activating the PLC and IP3R signaling pathway, which mediates Ca^2+^ release from the ER^26–28^. This shared signaling axis led us to hypothesize that many hit compounds might attenuate exodosis by antagonizing the IP3R pathway. Indeed, our activity-based classification showed that many hits associated with these GPCRs exerted broad inhibitory effects, potentially acting as pan-antagonists. For example, bromocriptine, a previously characterized hit^6^, has established clinical activities in the context of Parkinson’s and diabetes, but its relationship to ER stress and exodosis are incompletely understood. Our SCOPE analysis highlighted bromocriptine’s broad polypharmacology across several GPCRs, suggesting a more complex mechanism of action beyond its canonical dopamine D2 receptor agonism, and offering a plausible explanation for its anti-exodosis activity. Still, some hits displayed preferential inhibition of specific GPCRs, such as HRH1, HTR2A, DRD2, ADRA1A, HTR6, and SLC6A2.

We experimentally validated the functional relevance of one of these targets, HRH1, using selective small molecule antagonists and agonists. Selective antagonists of HRH1 (clemastine, rupatadine, and azelastine) consistently suppressed thapsigargin-induced secretion of GLuc SERCaMP, supporting a role for histaminergic signaling in exacerbating exodosis. In contrast, the HRH1 selective agonists (*N-*methylhistaprodifen and 2-pyridylethylamine) enhanced secretion, further strengthening the hypothesis that HRH1 activation potentiates ER calcium stress and exodosis. Furthermore, RNAi-mediated knockdown of HRH1 led to a significant reduction in SERCaMP signal without affecting cell viability, providing genetic confirmation of HRH1’s contribution to ER calcium dynamics and its role in mediating the exodosis phenotype. These findings are consistent with prior studies showing that HRH1 activity influences intracellular calcium homeostasis and ER stress^34–36^. Our results thus position HRH1 as a novel modulator and a potential therapeutic target in disorders associated with ER stress and exodosis.

Despite the significant advancements offered by SCOPE and the novel insights into exodosis, this study is not without limitations. First, while SCOPE integrates data from five major public databases, target association data could not be retrieved for 3,295 out of 10,124 compounds in the screening library, and 42 of the 185 hits lacked target annotations. This reflects the broader challenge of incomplete target annotation for many small molecules, especially investigational or less-characterized compounds, which may limit the scope of enrichment analyses. To improve coverage, future efforts could integrate additional resources or employ predictive modeling to infer targets for compounds without known annotations. Second, the experimental validation of HRH1’s role in ER exodosis was primarily conducted in SH-SY5Y human neuroblastoma cells. While this cell line is relevant for neurodegeneration models, the findings may not be fully generalizable to other cell types or in vivo systems, which could exhibit different ER stress responses, histamine receptor expression, or functionality of histamine receptor signaling pathways. Additionally, further experimental validation of the identified GPCR targets, beyond HRH1, is warranted to fully elucidate their roles in exodosis and ER calcium dynamics.

While demonstrated here in the context of ER proteostasis, we anticipate that SCOPE will be broadly applicable to other complex phenotypic screening campaigns. Applying the pipeline to screens across diverse biological processes and disease areas will further establish its versatility and utility in accelerating mechanism-agnostic drug discovery and drug repurposing.

In summary, SCOPE provides a robust and scalable computational framework for elucidating mechanisms of action from phenotypic screening data. Its application to ER proteome stabilizers not only revealed novel regulators of exodosis, such as HRH1, but also demonstrated the broader utility of compound-target enrichment strategies in deconvoluting polypharmacological effects. As phenotypic screening continues to grow in scale and complexity, tools like SCOPE will be critical for bridging high-throughput discovery efforts with systems-level interpretation, advancing our ability to identify new mechanisms, therapeutic targets, and drug repurposing opportunities.

## Online Methods

### Screening Compound Ontology for Pathway Enrichment (SCOPE) Pipeline

SCOPE is a computational pipeline that connects the pharmacologically diverse set of hits obtained in high-content screens (HCS) or high-throughput screens (HTS) to their biological targets and pathways providing hypothesis on mechanism of actions shared by hits. Fig. 1 provides a graphical overview of different steps involved in the SCOPE pipeline which is implemented as a KNIME^37^ workflow (version 4.5.3). Below we describe briefly each step of the pipeline:

#### Input data: using stabilizers of endoplasmic reticulum resident proteome as case study

The input of the workflow is a list of compounds, corresponding to the screening collection from which hits were derived, provided as an Excel file consisting of two columns named: ‘SMILES’ and ‘Sample ID’. Sample id is a compound identifier associated with the SMILES representation of the input compound and it can be any string (representing name or compound id). Another optional column named ‘Gene Symbol’ can be provided which lists a single gene name (such as a primary annotation) known to be associated with a compound (see more details under data sources for compound-target interactions section). A second input file is also required listing the ‘Sample ID’ of the compounds identified as hits. This file is used to select the subset of compound-target profiles for the hit compounds from the full set of compound-target profiles for the screening collection.

To demonstrate the utility of our workflow in interpreting hits resulting from a phenotypic screen we applied the workflow to compounds identified as stabilizers of the ER resident proteome^6^. Depletion of ER calcium can lead to the loss of resident proteins in a process termed exodosis. We previously performed a target-agnostic quantitative HTS using the GLuc SERCaMP assay, which monitors secretion of ER-resident proteins triggered by calcium depletion^6^. The assay was used to screen 10,124 compounds (9,501 unique compounds), including a collection of drugs approved by the FDA (US), EMA (EU), NHI (Japan), and HC (Canada) regulatory agencies. Compounds that decreased GLuc SERCaMP release, were subjected to counterscreens to filter out compounds that inhibited either secretory pathways, transgene expression, or cell viability resulting in the identification of 185 compounds that potentially stabilizes ER-resident proteome by different mechanisms. The 185 hits along with the screening collection of 10,124 compounds was used as input sets and subjected to the SCOPE workflow as described in the following sections.

#### Compound standardization

Input compounds were provided as SMILES strings and standardized using RDKit structure normalizer node which applies a series of transformations (removal of hydrogens, stripping of salts, recombine charges, etc.) to correct for any ambiguities in the input SMILES. The canonical SMILES of the corrected structure are then converted into InChIKey (IUPAC International Chemical Identifier Key) representation. For molecules containing salt, InChIKeys were generated both with and without salts to support flexible matching in downstream analyses.

#### Generating compound-target profiles

Target profiles for each input compound are built by querying five different databases: PubChem^22^, IUPHAR^21^, PharmGKB^23^, ChEMBL^20^, and DrugBank^24^. These databases either provide information on proven compound-target association (such as DrugBank and PharmGKB) or contain target-associated bioactivity (PubChem, IUPHAR, and ChEMBL). We only consider human proteins as targets of interest. Furthermore, we apply a 10 µM cutoff for bioactivity-based target annotations. The target annotations obtained from the five databases are optionally further supplemented with any gene annotation provided as input, adding a layer of flexibility to our workflow allowing the user to provide their own annotations to be considered for analysis. The metadata associated with target annotations from all resources are written out to CSV files. To query these databases, we first use the UniChem^38^ web API to map the compound InChIKey to corresponding compound identifiers in the databases. Below we describe briefly the steps followed to generate target profiles from different sources:

##### DrugBank

The DrugBank database (https://go.drugbank.com/) is a freely accessible comprehensive resource containing information about drugs and their targets. DrugBank provides access to its data in the form of XML file for noncommercial usage by obtaining the free academic license. Our workflow uses the drugbank id of the input compounds obtained through UniChem mapping to filter the XML file for relevant compounds and extract the targets information. Any non-human target proteins and those with ‘unknown’ under known-action attribute (which provides information on whether the pharmacological action of the drug is due to a specific target interaction) are filtered out.

##### PharmGKB

Pharmacogenomics Knowledgebase (PharmGKB) database (https://www.pharmgkb.org) contains data about genes, drugs, diseases, and the pharmacogenomic relationships between them. Relationship files were downloaded from the database and PharmGKB compound id mapped by UniChem was used to parse the file for compound-target associations. Pharos^39^ (https://pharos.nih.gov/), an interface to the Target Central Resource Database (TCRD; https://habanero.health.unm.edu/tcrd/) which curates information about human targets, was used to map the gene symbols retrieved from the relationship file to UniProt accession number.

##### IUPHAR

IUPHAR/BPS Guide to Pharmacology database (https://www.guidetopharmacology.org/) is an expert-curated resource of ligand-activity-target relationships and it offers access to its data as JSON files through web services. UniChem mapped IUPHAR id of the input compounds are first used to retrieve the interactions data from the database which is then parsed to remove activity data above 10 µM and those against non-human targets. The target ids obtained is further parsed to retrieve the gene symbol. The UniProt accession numbers of target genes are then queried via Pharos.

##### PubChem

PubChem (https://pubchem.ncbi.nlm.nih.gov/) is an open chemistry database that collects information from over 500 data sources providing access to various types of chemical information including compound biological activities^22^. PubChem Structured Data Query (SDQ) agent was used to retrieve structured-machine readable data by submitting queries as HTTP URLs. The UniChem mapped PubChem compound identifiers are used to generate a SDQ query to retrieve all the bioactivity data associated with human targets for each input compounds. The data is then parsed to retrieve target information removing any activity data above 10 µM.

##### ChEMBL

ChEMBL (https://www.ebi.ac.uk/chembl/) is a manually curated database of drug-like molecules with bioactivity data extracted from published literatures, deposited datasets, and patents^20^. Compound activity data from ChEMBL was accessed via MySQL queries to an instance of ChEMBL version 33.0. For each input compound, all human target associated bioactivity records corresponding to binding and functional assays (or patents) with a target type of single protein and a pChEMBL value of 5 or above are extracted. pChEMBL is defined as: −log10 (molar IC50, XC50, EC50, AC50, Ki, Kd or Potency).

##### Input gene symbol

Pharos is used to map any user-provided gene symbol associated with input compounds to their corresponding UniProt accession number. This part of the workflow is optional and will be ignored in case the user inputs a blank column under ‘Gene Symbol’. For our input, we used the primary target annotations from our in-house database.

#### Collating targets information from all sources

The compound-target profiles generated from different sources are combined, removing any duplicate pairs, and processed to generate three tables. The first output table contains a list of all unique targets along with columns indicating which sources the target annotation was derived from. This table is used as input to a python script node which constructs an upset plot providing an intuitive subset-level visualization of target annotations from different sources. The second table contains the compound-target pairs for all the compounds with resolved targets and is further used for enrichment analysis (see next section). This table is connected to python nodes that plot the distribution of targets per compound and compounds per target. The third table lists for each compound the number of targets it associated with from different sources and an upset plot summarizing the annotation sources of compounds is also generated. These tables and any unresolved compounds are written to CSV files.

#### Target enrichment analysis

Depending on the number of hit compounds being analyzed, the compound-target associations may retrieve several hundreds of targets. Gaining biological insight from such a long list of protein targets is a cumbersome task and may lead to investigator bias toward a hypothesis of interest. Moreover, not all targets in the list will be relevant to the underlying biology as the presence of a target in the hit list will be influenced by the distribution of targets for the entire screening collection (background targets) assayed. To identify targets that are enriched in the hit list compared to background we carry out a statistical overrepresentation analysis (ORA) that analyzes whether compound associations with certain targets are overrepresented in our hit list. For each target, a contingency table (shown in Table 1) was constructed, and ORA was done using Fisher’s exact test. The p-values resulting from Fisher’s test was adjusted for multiple comparisons using the Benjamini-Hochberg procedure^40^.

**Table 1:**
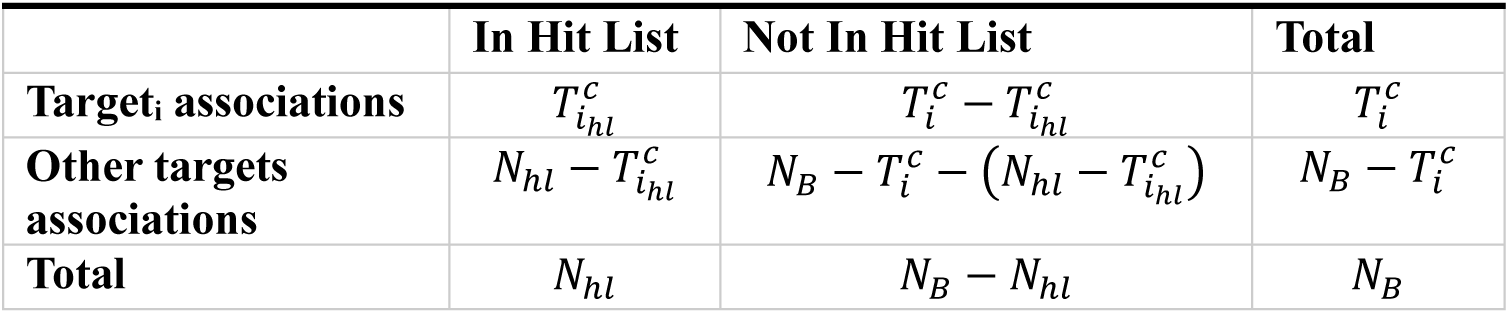
Contingency table for overrepresentation analysis. *N*_*B*_ is the number of compound-target associations for the entire screening collection (background targets), 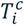 is the number of compound-associations for target_i_ in background, 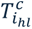 is the number of compound associations for target_i_ in the hit list, and *N*_*hl*_is the number of compound-target associations in the hit list.

#### Functional enrichment analysis

In the final step of the workflow, the enriched targets are queried through STRING database^25^ to fetch the protein-protein interaction networks as well as to connect targets to biological pathways.

### Activity based compound action classification

To classify the 55 hit compounds associated with the 26 enriched targets, we analyzed activity data from PubChem, ChEMBL, and IUPHAR. A compound was classified as an inhibitor if its reported activity against a given target included parameters such as *K_i_*, *pK_i_*, *INH*, or any inhibitory concentration values (e.g., IC_50_, IC_20_, IC_80_). Conversely, compounds with reported *EC* values (e.g., EC_50_, EC_20_) were classified as activators. Because *K_i_* and *pK_i_* values do not always reflect inhibitory action, we cross-referenced the ‘Action’ field from IUPHAR and the ‘Action Type’ field from ChEMBL, both of which indicate mechanism of action, to correct potential misclassifications when this information was available. If the reported activity metric was unrelated to activation or inhibition (e.g., *K_d_*, *K_b_*, *K_m_*, *K_a_*), the compound was classified as inconclusive. Compounds belonging to both activators and inhibitors for the same target were labeled as having a mixed annotation. Finally, if a compound-target association originated from curated databases such as DrugBank or PharmGKB without specific activity parameters, it was labeled as unknown.

### SERCaMP assay

Stable SH-SY5Y cells expressing the GLuc SERCaMP^6^ were grown in 10% serum-containing DMEM based media, supplemented with 1% Penn/Strep. Cells were plated at a density of 5×10^4^ cells/well in a 96-well tissue culture treated plate. After 24 hours, the media was exchanged for 1.5% serum-containing media. 24 hours after that, designated T0, the cells were treated with 10 μM H1R antagonists or agonists. At 16 hours, the cultures that were treated with the histamine antagonists were treated with 100 nM thapsigargin for an additional 4 hours. 10 μL supernatant samples were collected at T16 and T20 hour timepoints and transferred to an opaque-walled plate for luciferase activity evaluation. 100 μL of a 10 μM coelenterazine (Regis Technologies, Morton Grove, IL) in PBS solution was injected into each well by a BioTek Synergy Neo2 Hybrid Multimode Reader (Biotek Synergy II, Winooski, VT) and luminescence subsequently read.

### RNAseq

SH-SY5Y cells were plated at a concentration of 3.12×10^5^ cells/well in a 24 well plate. Cells were treated with 0.1% dimethyl sulfoxide (DMSO) for 30 min followed by an additional 0.1% DMSO treatment for 8 hrs. RNA was harvested and RNA integrity was evaluated by RIN measurement. RNA samples were sent to Novogene to perform bulk RNA sequencing and analysis. Expression of genes: HRH1 (ENSG00000196639), HRH2 (ENSG00000113749), HRH3 (ENSG00000101180), HRH4 (ENSG00000134489) were assessed.

### RNAi

siRNA complexes were formed in 384-well opaque tissue culture treated plates by combining 2 μL of 20 nM siRNA with 20 μL of transfection solution containing 0.15 μL Lipofectamine RNAiMAX (Thermo) in 20uL of DMEM + 1 mM sodium pyruvate. siRNA sequences for HRH1 were s6902 5’-GUAUCUGGGUUGCACAUGAtt-3’; s223879 5’-GGCAAAGGCAAAUUGAGGAtt-3’; and s6904 5’-UCCUGUGCAUUGAUCGCUAtt-3’. The mixture was incubated at room temperature for 30 min to allow complexes to form. SH-SY5Y cells expressing SERCaMP were suspended in DMEM + 1 mM sodium pyruvate + 4 mM L-glutamine + 20% FBS + 200 U/mL penicillin, 200 μg/mL streptomycin, to a density of 1×10^5^ cells/mL and 20 μL of the cell suspension (2000 cells) was added to the 384-well plate containing siRNA complexes. Cells were incubated at 37C, 95% RH, 5% CO_2_ for 96 h. Thapsigargin was prepared at 1.5 μM concentration in DMEM + 1 mM sodium pyruvate + 2 mM L-glutamine + 10% FBS + 100 U/mL penicillin, 100 μg/mL streptomycin and 10 μl was added to the well (300 nM final concentration of thapsigargin). Cells were incubated at 37C, 95% RH, 5% CO2 for 4h. 5 μL of the medium was transferred to a new opaque plate containing 20 μL of PBS and the secreted GLuc-SERCaMP was measured by adding 5 μL of Pierce Gaussia Luciferase Glow Assay Reagent, prepared according to manufacturer’s instructions. Luminescence was measured using an Envision Multimode Detection system (Revvity) equipped for luminescence. Viability was measured in the original plate containing cells by adding 20 μl of CellTiterGlo Reagent (Promega) and measuring luminescence on the Envision. Median values were calculated for the SERCaMP and viability assays.

## Data availability statement

The KNIME workflow developed and used in this study will be made publicly available on the KNIME Community Hub upon acceptance of the manuscript.

## Acknowledgments

We thank Dr. Madhu Lal and the RNAi Screening Facility at the National Center for Advancing Translational Sciences for providing data on RNAi screen. We thank Dr. Emily Wires for technical consultation and Lana Gore and Lacey K. Greer for technical assistance. This research was supported by the intramural research programs at the National Institute on Drug Abuse and the National Center for Advancing Translational Sciences, National Institutes of Health.

## NNotes

The authors declare no competing financial interest.

## References

1 Vincent, F. et al. Phenotypic drug discovery: recent successes, lessons learned and new directions. Nat Rev Drug Discov 21, 899–914 (2022). 10.1038/s41573-022-00472-w

2 Cooper, D. J., Zunino, G., Bixby, J. L. & Lemmon, V. P. Phenotypic screening with primary neurons to identify drug targets for regeneration and degeneration. Mol Cell Neurosci 80, 161–169 (2017). 10.1016/j.mcn.2016.07.001

3 Moffat, J. G., Vincent, F., Lee, J. A., Eder, J. & Prunotto, M. Opportunities and challenges in phenotypic drug discovery: an industry perspective. Nat Rev Drug Discov 16, 531–543 (2017). 10.1038/nrd.2017.111

4 Ryoo, H., Kimmel, H., Rondo, E. & Underhill, G. H. Advances in high throughput cell culture technologies for therapeutic screening and biological discovery applications. Bioeng Transl Med 9, e10627 (2024). 10.1002/btm2.10627

5 Mekahli, D., Bultynck, G., Parys, J. B., De Smedt, H. & Missiaen, L. Endoplasmic-reticulum calcium depletion and disease. Cold Spring Harb Perspect Biol 3 (2011). 10.1101/cshperspect.a004317

6 Henderson, M. J. et al. A target-agnostic screen identifies approved drugs to stabilize the endoplasmic reticulum-resident proteome. Cell Rep 35, 109040 (2021). 10.1016/j.celrep.2021.109040

7 Martinez, G. et al. Regulation of Memory Formation by the Transcription Factor XBP1. Cell Rep 14, 1382–1394 (2016). 10.1016/j.celrep.2016.01.028

8 Plate, L. et al. Small molecule proteostasis regulators that reprogram the ER to reduce extracellular protein aggregation. Elife 5 (2016). 10.7554/eLife.15550

9 Zou, H. et al. Benzodiazepinone derivatives protect against endoplasmic reticulum stress-mediated cell death in human neuronal cell lines. ACS Chem Neurosci 6, 464–475 (2015). 10.1021/cn500297v

10 Trychta, K. A., Back, S., Henderson, M. J. & Harvey, B. K. KDEL Receptors Are Differentially Regulated to Maintain the ER Proteome under Calcium Deficiency. Cell Rep 25, 1829–1840 e1826 (2018). 10.1016/j.celrep.2018.10.055

11 Xu, C., Bailly-Maitre, B. & Reed, J. C. Endoplasmic reticulum stress: cell life and death decisions. J Clin Invest 115, 2656–2664 (2005). 10.1172/JCI26373

12 Trychta, K. A. & Harvey, B. K. Caffeine and MDMA (Ecstasy) Exacerbate ER Stress Triggered by Hyperthermia. Int J Mol Sci 23 (2022). 10.3390/ijms23041974

13 Dossat, A. M. et al. Excitotoxic glutamate levels cause the secretion of resident endoplasmic reticulum proteins. J Neurochem 168, 2461–2478 (2024). 10.1111/jnc.16093

14 Napolitano, F., Sirci, F., Carrella, D. & di Bernardo, D. Drug-set enrichment analysis: a novel tool to investigate drug mode of action. Bioinformatics 32, 235–241 (2016). 10.1093/bioinformatics/btv536

15 Huang, C. et al. The DrugPattern tool for drug set enrichment analysis and its prediction for beneficial effects of oxLDL on type 2 diabetes. J Genet Genomics 45, 389–397 (2018). 10.1016/j.jgg.2018.07.002

16 Kropiwnicki, E. et al. Drugmonizome and Drugmonizome-ML: integration and abstraction of small molecule attributes for drug enrichment analysis and machine learning. Database (Oxford) 2021 (2021). 10.1093/database/baab017

17 Park, S. et al. CSgator: an integrated web platform for compound set analysis. J Cheminform 11, 17 (2019). 10.1186/s13321-019-0339-6

18 Fuzi, B., Gurinova, J., Hermjakob, H., Ecker, G. F. & Sheriff, R. Path4Drug: Data Science Workflow for Identification of Tissue-Specific Biological Pathways Modulated by Toxic Drugs. Front Pharmacol 12, 708296 (2021). 10.3389/fphar.2021.708296

19 Jassal, B. et al. The reactome pathway knowledgebase. Nucleic Acids Res 48, D498–D503 (2020). 10.1093/nar/gkz1031

20 Zdrazil, B. et al. The ChEMBL Database in 2023: a drug discovery platform spanning multiple bioactivity data types and time periods. Nucleic Acids Res 52, D1180–D1192 (2024). 10.1093/nar/gkad1004

21 Harding, S. D. et al. The IUPHAR/BPS Guide to PHARMACOLOGY in 2024. Nucleic Acids Res 52, D1438–D1449 (2024). 10.1093/nar/gkad944

22 Kim, S. et al. PubChem 2023 update. Nucleic Acids Res 51, D1373–D1380 (2023). 10.1093/nar/gkac956

23 Whirl-Carrillo, M. et al. An Evidence-Based Framework for Evaluating Pharmacogenomics Knowledge for Personalized Medicine. Clin Pharmacol Ther 110, 563–572 (2021). 10.1002/cpt.2350

24 Knox, C. et al. DrugBank 6.0: the DrugBank Knowledgebase for 2024. Nucleic Acids Res 52, D1265–D1275 (2024). 10.1093/nar/gkad976

25 Szklarczyk, D. et al. The STRING database in 2023: protein-protein association networks and functional enrichment analyses for any sequenced genome of interest. Nucleic Acids Res 51, D638–D646 (2023). 10.1093/nar/gkac1000

26 Mahmood, A., Ahmed, K. & Zhang, Y. beta-Adrenergic Receptor Desensitization/Down-Regulation in Heart Failure: A Friend or Foe? Front Cardiovasc Med 9, 925692 (2022). 10.3389/fcvm.2022.925692

27 Okubo, Y. Astrocytic Ca(2+) signaling mediated by the endoplasmic reticulum in health and disease. J Pharmacol Sci 144, 83–88 (2020). 10.1016/j.jphs.2020.07.006

28 Selway, J. L., Moore, C. E., Mistry, R., John Challiss, R. A. & Herbert, T. P. Molecular mechanisms of muscarinic acetylcholine receptor-stimulated increase in cytosolic free Ca(2+) concentration and ERK1/2 activation in the MIN6 pancreatic beta-cell line. Acta Diabetol 49, 277–289 (2012). 10.1007/s00592-011-0314-9

29 Shamizadeh, S., Brockow, K. & Ring, J. Rupatadine: efficacy and safety of a non-sedating antihistamine with PAF-antagonist effects. Allergo J Int 23, 87–95 (2014). 10.1007/s40629-014-0011-7

30 Corren, J. et al. Effectiveness of azelastine nasal spray compared with oral cetirizine in patients with seasonal allergic rhinitis. Clin Ther 27, 543–553 (2005). 10.1016/j.clinthera.2005.04.012

31 Dorsch, W., Reimann, H. J. & Neuhauser, J. Histamine1--histamine2 antagonism: effect of combined clemastine and cimetidine pretreatment on allergen and histamine-induced reactions of the guinea pig lung in vivo and in vitro. Agents Actions 12, 113–118 (1982). 10.1007/BF01965120

32 Seifert, R. et al. Multiple differences in agonist and antagonist pharmacology between human and guinea pig histamine H1-receptor. J Pharmacol Exp Ther 305, 1104–1115 (2003). 10.1124/jpet.103.049619

33 Ratnala, V. R. et al. Large-scale overproduction, functional purification and ligand affinities of the His-tagged human histamine H1 receptor. Eur J Biochem 271, 2636–2646 (2004). 10.1111/j.1432-1033.2004.04192.x

34 Chen, W. et al. Maprotiline restores ER homeostasis and rescues neurodegeneration via Histamine Receptor H1 inhibition in retinal ganglion cells. Nat Commun 13, 6796 (2022). 10.1038/s41467-022-34682-y

35 Fan, J. et al. Modulating endoplasmic reticulum stress attenuates mast cell degranulation. Int Immunopharmacol 126, 111336 (2024). 10.1016/j.intimp.2023.111336

36 Deniaud, A. et al. Endoplasmic reticulum stress induces calcium-dependent permeability transition, mitochondrial outer membrane permeabilization and apoptosis. Oncogene 27, 285–299 (2008). 10.1038/sj.onc.1210638

37 Berthold, M. R., et al. KNIME:: The Konstanz Information Miner. Stud Class Data Anal, 319–326 (2008). Doi 10.1145/1656274.1656280

38 Chambers, J. et al. UniChem: a unified chemical structure cross-referencing and identifier tracking system. J Cheminform 5, 3 (2013). 10.1186/1758-2946-5-3

39 Kelleher, K. J. et al. Pharos 2023: an integrated resource for the understudied human proteome. Nucleic Acids Res 51, D1405–D1416 (2023). 10.1093/nar/gkac1033

40 Yekutieli, D. & Benjamini, Y. Resampling-based false discovery rate controlling multiple test procedures for correlated test statistics. J Stat Plan Infer 82, 171–196 (1999). Doi 10.1016/S0378-3758(99)00041-5

