## Supplemental Tables for "SCOPE: Revealing Hidden Mechanisms in Phenotypic Screens Through Target and Pathway Enrichment"

***Supplementary Table 1.*** List of enriched targets after target enrichment analysis. All targets with a 5% cutoff for FDR and p-value are listed.

| **Uniprot ID** | **Symbol** | **Protein Name** | **Fisher_p-value** | **FDR (BH)** |
|---|---|---|---|---|
| P28223 | HTR2A | 5-hydroxytryptamine receptor 2A | 1.58E-10 | 6.20E-08 |
| P28335 | HTR2C | 5-hydroxytryptamine receptor 2C | 2.25E-07 | 3.68E-05 |
| P08912 | CHRM5 | Muscarinic acetylcholine receptor M5 | 2.82E-07 | 3.68E-05 |
| P35348 | ADRA1A | Alpha-1A adrenergic receptor | 4.86E-07 | 4.76E-05 |
| P08172 | CHRM2 | Muscarinic acetylcholine receptor M2 | 8.27E-07 | 6.48E-05 |
| P35462 | DRD3 | D(3) dopamine receptor | 1.30E-06 | 8.52E-05 |
| P21728 | DRD1 | D(1A) dopamine receptor | 3.30E-06 | 1.85E-04 |
| P20309 | CHRM3 | Muscarinic acetylcholine receptor M3 | 4.59E-06 | 2.25E-04 |
| P50406 | HTR6 | 5-hydroxytryptamine receptor 6 | 6.91E-06 | 3.01E-04 |
| Q12809 | KCNH2 | Potassium voltage-gated channel subfamily H member 2 | 7.73E-06 | 3.03E-04 |
| P08173 | CHRM4 | Muscarinic acetylcholine receptor M4 | 2.29E-05 | 8.18E-04 |
| P14416 | DRD2 | D(2) dopamine receptor | 6.35E-05 | 1.92E-03 |
| P25100 | ADRA1D | Alpha-1D adrenergic receptor | 6.17E-05 | 1.92E-03 |
| P41595 | HTR2B | 5-hydroxytryptamine receptor 2B | 7.63E-05 | 2.14E-03 |
| P08908 | HTR1A | 5-hydroxytryptamine receptor 1A | 1.45E-04 | 3.54E-03 |
| P11229 | CHRM1 | Muscarinic acetylcholine receptor M1 | 1.37E-04 | 3.54E-03 |
| P23975 | SLC6A2 | Sodium-dependent noradrenaline transporter | 1.79E-04 | 3.89E-03 |
| P08913 | ADRA2A | Alpha-2A adrenergic receptor | 1.78E-04 | 3.89E-03 |
| P18089 | ADRA2B | Alpha-2B adrenergic receptor | 2.19E-04 | 4.52E-03 |
| P25021 | HRH2 | Histamine H2 receptor | 2.62E-04 | 4.89E-03 |
| P35372 | OPRM1 | Mu-type opioid receptor | 2.61E-04 | 4.89E-03 |
| P35367 | HRH1 | Histamine H1 receptor | 3.23E-04 | 5.51E-03 |
| P31645 | SLC6A4 | Sodium-dependent serotonin transporter | 3.11E-04 | 5.51E-03 |
| P46098 | HTR3A | 5-hydroxytryptamine receptor 3A | 7.44E-04 | 1.22E-02 |
| P21918 | DRD5 | D(1B) dopamine receptor | 1.43E-03 | 2.25E-02 |
| P41145 | OPRK1 | Kappa-type opioid receptor | 3.12E-03 | 4.71E-02 |
| P18825 | ADRA2C | Alpha-2C adrenergic receptor | 3.65E-03 | 5.29E-02 |

| Uniprot ID | Symbol | Protein Name | Fisher_p-value | FDR (BH) |
|---|---|---|---|---|
| P21917 | DRD4 | D(4) dopamine receptor | 4.42E-03 | 6.18E-02 |
| Q99720 | SIGMAR1 | Sigma non-opioid intracellular receptor 1 | 4.70E-03 | 6.35E-02 |
| Q9NY46 | SCN3A | Sodium channel protein type 3 subunit alpha | 5.72E-03 | 7.47E-02 |
| P08183 | ABCB1 | Multidrug resistance protein 1 | 6.88E-03 | 8.70E-02 |
| Q99250 | SCN2A | Sodium channel protein type 2 subunit alpha | 8.72E-03 | 9.76E-02 |
| P35498 | SCN1A | Sodium channel protein type 1 subunit alpha | 8.04E-03 | 9.76E-02 |
| Q14147 | DHX34 | Probable ATP-dependent RNA helicase DHX34 | 8.70E-03 | 9.76E-02 |
| Q14562 | DHX8 | ATP-dependent RNA helicase DHX8 | 8.70E-03 | 9.76E-02 |
| P47898 | HTR5A | 5-hydroxytryptamine receptor 5A | 1.29E-02 | 1.41E-01 |
| P04156 | PRNP | Major prion protein | 1.68E-02 | 1.73E-01 |
| Q7Z2K8 | GPRIN1 | G protein-regulated inducer of neurite outgrowth 1 | 1.68E-02 | 1.73E-01 |
| P34969 | HTR7 | 5-hydroxytryptamine receptor 7 | 2.15E-02 | 1.74E-01 |
| P25103 | TACR1 | Substance-P receptor | 2.34E-02 | 1.74E-01 |
| Q08499 | PDE4D | cAMP-specific 3',5'-cyclic phosphodiesterase 4D | 2.10E-02 | 1.74E-01 |
| Q16769 | QPCT | Glutaminyl-peptide cyclotransferase | 2.97E-02 | 1.74E-01 |
| Q86TP1 | PRUNE1 | Exopolyphosphatase PRUNE1 | 2.97E-02 | 1.74E-01 |
| P20648 | ATP4A | Potassium-transporting ATPase alpha chain 1 | 2.20E-02 | 1.74E-01 |
| Q05586 | GRIN1 | Glutamate receptor ionotropic, NMDA 1 | 1.96E-02 | 1.74E-01 |
| P28907 | CD38 | ADP-ribosyl cyclase/cyclic ADP-ribose hydrolase 1 | 2.97E-02 | 1.74E-01 |
| Q9Y5S1 | TRPV2 | Transient receptor potential cation channel subfamily V member 2 | 2.97E-02 | 1.74E-01 |
| P51878 | CASP5 | Caspase-5 | 2.97E-02 | 1.74E-01 |
| P49662 | CASP4 | Caspase-4 | 2.97E-02 | 1.74E-01 |
| P62937 | PPIA | Peptidyl-prolyl cis-trans isomerase A | 2.97E-02 | 1.74E-01 |
| Q08752 | PPID | Peptidyl-prolyl cis-trans isomerase D | 2.97E-02 | 1.74E-01 |
| P23284 | PPIB | Peptidyl-prolyl cis-trans isomerase B | 2.97E-02 | 1.74E-01 |
| P20815 | CYP3A5 | Cytochrome P450 3A5 | 2.04E-02 | 1.74E-01 |

| Uniprot ID | Symbol | Protein Name | Fisher_p-value | FDR (BH) |
|---|---|---|---|---|
| O14939 | PLD2 | Phospholipase D2 | 2.97E-02 | 1.74E-01 |
| Q13393 | PLD1 | Phospholipase D1 | 2.97E-02 | 1.74E-01 |
| P10909 | CLU | Clusterin | 2.97E-02 | 1.74E-01 |
| Q96LB1 | MRGPRX2 | Mas-related G-protein coupled receptor member X2 | 2.97E-02 | 1.74E-01 |
| Q13639 | HTR4 | 5-hydroxytryptamine receptor 4 | 2.65E-02 | 1.74E-01 |
| P49069 | CAMLG | Calcium signal-modulating cyclophilin ligand | 2.97E-02 | 1.74E-01 |
| Q96LZ3 | PPP3R2 | Calcineurin subunit B type 2 | 2.97E-02 | 1.74E-01 |
| P30405 | PPIF | Peptidyl-prolyl cis-trans isomerase F, mitochondrial | 2.97E-02 | 1.74E-01 |
| Q9H4I9 | SMDT1 | Essential MCU regulator, mitochondrial | 2.97E-02 | 1.74E-01 |
| Q9HAW9 | UGT1A8 | UDP-glucuronosyltransferase 1-8 | 2.20E-02 | 1.74E-01 |
| Q6ICB4 | PHETA2 | Sesquipedalian-2 | 2.97E-02 | 1.74E-01 |
| Q2M1K9 | ZNF423 | Zinc finger protein 423 | 2.97E-02 | 1.74E-01 |
| Q9NZI2 | KCNIP1 | Kv channel-interacting protein 1 | 2.97E-02 | 1.74E-01 |
| Q01726 | MC1R | Melanocyte-stimulating hormone receptor | 2.97E-02 | 1.74E-01 |
| Q92847 | GHSR | Growth hormone secretagogue receptor type 1 | 4.10E-02 | 2.36E-01 |
| P07550 | ADRB2 | Beta-2 adrenergic receptor | 4.74E-02 | 2.68E-01 |
| Q15125 | EBP | 3-beta-hydroxysteroid-Delta(8),Delta(7)-isomerase | 4.78E-02 | 2.68E-01 |

***Supplementary Table 2.*** Biological activity of bromocriptine against different human receptors.

Data taken from IUPHAR (https://www.guidetopharmacology.org/GRAC/LigandDisplayForward?tab=biology&ligandId=35).

| **Target** | **Action** | **Value** | **Parameter** |
|---|---|---|---|
| ADRA2A | Antagonist | 8.0 – 8.3 | $pK_i$ |
| HTR1D | Partial agonist | 8 | $pK_i$ |
| HTR1A | Partial agonist | 7.9 | $pK_i$ |
| DRD2 | Full agonist | 7.3 – 8.3 | $pK_i$ |
| DRD3 | Partial agonist | 7.1 – 8.2 | $pK_i$ |
| ADRA2C | Antagonist | 7.6 | $pK_i$ |
| HTR6 | Full agonist | 7.5 | $pK_i$ |
| HTR2B | Antagonist | 7.3 | $pK_i$ |
| ADRA2B | Antagonist | 6.9 – 7.5 | $pK_i$ |
| HTR2A | Partial agonist | 7 | $pK_i$ |
| HTR1B | Partial agonist | 6.5 | $pK_i$ |
| DRD4 | Antagonist | 6.4 | $pK_i$ |
| DRD5 | Full agonist | 6.3 | $pK_i$ |
| DRD1 | Partial agonist | 6.2 | $pK_i$ |
| HTR2C | Partial agonist | 6.1 | $pK_i$ |
